# Novel 3-D macrophage spheroid model reveals reciprocal regulation of immunomechanical stress and mechano-immunological response

**DOI:** 10.1101/2024.02.14.580327

**Authors:** Alice Burchett, Saeed Siri, Jun Li, Xin Lu, Meenal Datta

## Abstract

**Purpose:** In many diseases, an overabundance of macrophages contributes to adverse outcomes. While numerous studies have compared macrophage phenotype after mechanical stimulation or with varying local stiffness, it is unclear if and how macrophages themselves contribute to mechanical forces in their microenvironment.

**Methods:** Raw 264.7 murine macrophages were embedded in a confining agarose gel, where they proliferated to form spheroids over time. Gels were synthesized at various concentrations to tune the stiffness and treated with various growth supplements to promote macrophage polarization. The spheroids were then analyzed by immunofluorescent staining and qPCR for markers of proliferation, mechanosensory channels, and polarization. Finally, spheroid geometries were used to computationally model the strain generated in the agarose by macrophage spheroid growth.

**Results:** Macrophages form spheroids and generate growth-induced mechanical forces (i.e., solid stress) within confining agarose gels, which can be maintained for at least 16 days in culture. Increasing agarose concentration restricts spheroid expansion, promotes discoid geometries, limits gel deformation, and induces an increase in iNOS expression. LPS stimulation increases spheroid growth, though this effect is reversed with the addition of IFN-γ. Ki67 expression decreases with increasing agarose concentration, in line with the growth measurements.

**Conclusions:** Macrophages alone both respond to and generate solid stress. Understanding how macrophage generation of growth-induced solid stress responds to different environmental conditions will help to inform treatment strategies for the plethora of diseases that involve macrophage accumulation.

## Introduction

From atherosclerotic plaques to solid cancerous tumors, macrophages play important roles in immune effector function and orchestration, and as active constituents of the mechanical microenvironment [1]. As innate immune cells, they are not only part of the first line of defense against pathogens, they also contribute to tissue repair and help coordinate the broader immune response. Macrophage phenotype is highly plastic and ebbs between pro- and anti-inflammatory states, as these cells sense and correspondingly respond to diverse and dynamic microenvironments [2]. Macrophages present in tissues are either resident or derived from circulating monocytes that differentiate into macrophages upon vascular extravasation and tissue penetration [3]. During an inflammatory response, an injured or diseased site will accumulate macrophages, both through the recruitment of circulating monocytes and local proliferation of bone marrow and embryonic-derived macrophages [4]. For example, upon tissue injury, vascular endothelial cells upregulate adhesion molecules that allow patrolling monocytes to adhere to the vessel wall, where they withstand shear stress from blood flow, and eventually enter the tissue between endothelial junctions [5].

While macrophage phenotype and function, known also as polarization, is traditionally thought to be either pro-inflammatory (“M1-like”) or anti-inflammatory (“M2-like”), their dynamic cell state can lie on a continuum between inflammation-accelerating and inflammation-inhibiting responses [6]. Macrophages adopt and shift between these polarizations in any tissue; generally, equilibrium between the two ends of the spectrum is essential for homeostatic tissue maintenance and repair. However, an over- or under-active macrophage response can disrupt this tenuous balance, particularly in cases where macrophages accumulate in large quantities.

Elevated macrophage populations in diseased tissue are often correlated with worse prognosis, particularly in diseases where altered tissue mechanics contribute to the pathophysiology [7]. Cancer, wound healing, bacterial infections, and other disease settings involve altered tissue mechanics, which impact immune surveillance [8]. For example, atherosclerotic plaques physically disrupt blood flow, and their mechanical stability determines if they will rupture, leading to downstream ischemic events [9]. Macrophages accumulate in plaques, where they contribute to this mechanical instability, increasing the risk of life-threatening events such as a stroke. Macrophages infiltrate tumor microenvironments in high numbers, with the density of tumor-associated macrophages correlating with worse prognoses in cancers including glioblastoma, ovarian cancer, and breast cancer [10–14]. In glioblastoma, macrophages can comprise up to 50% of the tumor bulk, promoting tumor progression and treatment resistance [15–17]. Macrophages also contribute to increased collagen deposition in hypertrophic scars and heart attack scarring, leading to diminished tissue function [18, 19].

In addition to classical biochemical cues, macrophages also respond to mechanical stimuli such as shear stress, tissue viscoelasticity, cyclic compression or stretching, and hydrodynamic pressure changes [20]. This response has been characterized in prior studies, particularly in the context of the cardiovascular and skeletomuscular systems [21–24]. Macrophages experience a wide range of tissue mechanical properties, with Young’s moduli on the order of single kilopascals in the brain to tens of gigapascals in bone [25, 26]. *In vitro* experiments show that macrophages have a stronger inflammatory response when cultured on stiffer 2-D substrates [27–31]. However, the opposite effect is observed when cells are cultured in a 3-D matrix. Macrophages in stiffer matrices *in vitro* and *in vivo* have a more immunosuppressive, M2-like phenotype [32–34]. Thus, a physiologically relevant *in vitro* model is essential to understanding how macrophages respond – and contribute – to mechanical stimuli in the body.

Studies on macrophage mechanical responses largely neglect the impact of growth-induced mechanical forces – including solid stress [35], generated by solid tissue components (cells and matrix) – on macrophage proliferation. Further, the degree to which macrophages directly contribute to mechanical stress through their physical presence, accumulation, and proliferation in tissue remains unexplored. Here, we aimed to characterize the solid stress that macrophages generate through 3-D growth in a confining agarose gel, simulating the mechanics of the tissue microenvironment independent of confounding biochemical cues or matrix degradation.

Macrophages (RAW264.7) were embedded in agarose gels of varying substrate concentrations to span a range of physiologically-relevant stiffnesses. As the agarose-embedded cells proliferated to form spheroids, they displaced the surrounding gel, in a similar manner to a cancerous tumor generating and exerting solid stress on the surrounding tissue [36]. Spheroids in softer gels reached much larger sizes and caused larger displacements of the surrounding gel compared to spheroids in stiffer gels. Pro-inflammatory stimulation with lipopolysaccharide (LPS) also led to an increase in spheroid size, though this effect was reversed with the addition of IFNγ. The mechanosensitive ion channels Piezo1 and transient receptor potential vanilloid 4 (TRPV4) both decreased in trend with increased stiffness. Markers of both pro- and anti-inflammatory functional status both showed a trending increase with stiffness, though only the pro-inflammatory marker reached statistical significance. Overall, this work highlights a novel, tunable, and high throughput method of interrogating macrophage immunomechanics and mechano-immunology, with relevance to a wide range of diseases.

## Materials and Methods

### Cell culture, gel formation, and macrophage activation/polarization

RAW264.7 murine macrophages were purchased from ATCC (TIB-71). They were grown in a complete culture medium consisting of DMEM (Corning, 10-013-CV) supplemented with 10% Fetal Bovine Serum (Gibco, 26140079) and 1% penicillin-streptomycin (Corning, 30-002-CI). They were maintained in a humidified incubator at 37°C with 5% CO_2_. Cells were grown as adherent monolayers and passaged using a cell scraper for mechanical detachment.

To create agarose-embedded 3-D cultures, single-cell suspensions were mixed with low-gelling temperature agarose (Sigma-Aldrich, A0701-25G). First, a 4% agarose solution was made in a complete culture medium and heated in a microwave until dissolved. The liquid agarose was maintained at 48°C until use. Single-cell suspensions of RAW264.7 cells in medium were made ranging between 1000 cells/mL to 10,000,000 cells/mL and mixed with a proportional amount of the 4% agarose to create gels of 0.5%, 1%, or 2% agarose. The cell-agarose solution was pipetted into 2 mm-deep cylindrical molds and left at room temperature to set for 10 minutes. The gels were then removed from the molds, submerged in culture medium, and maintained under standard culture conditions.

For macrophage-stimulation-treated gels, the medium was supplemented with 200 ng/ml LPS. For M1-polarization-treated gels, the medium was supplemented with 20 ng/ml IFN-γ (BioLegend, 575302) and 200 ng/ml LPS (Santa Cruz Biotechnology, Inc, sc-3535). For M2-polarization-treated gels, the medium was supplemented with 20 ng/ml IL-4 (Pepro-Tech, 214-14). The volume of the gel was included in the total solution volume to achieve the final concentrations. Gels cultured under hypoxic conditions were placed in a Tri-Gas hypoxia incubator with 5% CO_2_ and 1% O_2_. Compressed gels were placed on a 0.4 μm pore size transwell cell culture insert (CellQART, 9310402) and a 3-D printed PLA weight was placed on top to apply 0.15 kPa of compression to the gel to simulate the solid stress measured in murine glioblastoma models [17, 37, 38]. A 1% agarose cushion was placed between the cells and the weight to serve as a media reservoir and protect cells from direct contact with the rigid weight.

### Staining whole agarose gels

For live/dead staining, the gels were incubated with 2 μg/ml Calcein-AM (BioLegend, 425201) and 1 μg/ml Propidium Iodide (MP Biomedicals, 0219545810) in complete medium for 30 minutes at 37°C. These were imaged on a point-scanning confocal microscope (Nikon AXR). To obtain 3-D images of the spheroids for strain quantification, the gels were fixed overnight in 4% paraformaldehyde in PBS (Thermo Fisher, J19943.K2) at 4°C, then rinsed with PBS and incubated with CellMask™ Orange Actin Tracking Stain (1:1000, Thermo Fisher, A57244) and DAPI (2 μg/ml Sigma-Aldrich, D9542) for 48 hours at 4°C. Image stacks were acquired using a multiphoton microscope (Leica Stellaris 8 DIVE).

### Immunofluorescent staining

After fixation as described above, gels were immersed in 30% sucrose overnight at 4°C, then transferred to a 1:1 solution of 30% sucrose and OCT (VWR, 95057-838) overnight at 4°C. The gels were then frozen in OCT and sectioned into 5 μm thick slices. The sections were incubated with primary antibodies for Ki67 (1:100, Novus Biologicals, NB110-89719) or Piezo1 (1:100, Proteintech, 15939-1-AP) overnight at 4°C. The following day, the slides were incubated with an anti-rabbit secondary antibody (1:500, abcam, ab150077) and DAPI and imaged (Leica DMi8).

### Image analysis

Spheroid contour plots were generated in Autodesk Inventor using surface files generated by ImageJ. For size comparison, spheroids were identified manually from brightfield images and analyzed using ImageJ’s Analyze Particles function. For immunofluorescent-stained sections, the spheroid area was identified by CellProfiler using the combined antibody and DAPI channels, and the average intensity within the spheroid area was recorded.

### PCR

RNA was collected from agarose-embedded cells after 48 hours in culture. For each sample, a small piece of gel (∼100 μl volume) was dissolved in 400 μl TRI-reagent (Zymo Research, R2050-1-200) and the RNA was purified using an RNA isolation kit (Zymo Research, R2051). Gene expression was analyzed using TaqMan primers for Piezo1 (Mm01241549_m1), TRPV4 (Mm00499025_m1), Ki67 (Mm01278617_m1), Caspase 3 (Mm01195085_m1), and GAPDH (Mm99999915_g1). Gene expression was normalized to GAPDH reported as 2^− ΔΔCt^.

### Computational modeling

To investigate the distribution of solid stress around the spheroids, a computational modeling approach was employed using COMSOL Multiphysics. Agarose gels with concentrations of 0.5%, 1%, and 2% were modeled as linear elastic materials (E = 2000, 19830, and 99596 Pa respectively). The spheroid geometry was obtained through image processing techniques and subsequently implemented in COMSOL. The initial geometry of the spheroid was considered as a sphere with a 5-micrometer radius. Through the application of prescribed displacement boundary conditions, the spheroids were expanded to their final geometry within the agarose gel. This approach allowed for the exploration of the solid stress and displacement distribution around the spheroids. The modeling simulations were conducted in 2D, providing a comprehensive analysis of the mechanical stress distribution within the agarose gel environment. This methodological approach enables a detailed examination of the impact of agarose gel concentration on the mechanical behavior of macrophage spheroids in a controlled and reproducible manner.

### Statistical Analysis

Statistical analyses and data visualization were done using Graphpad Prism. For spheroid viability and size analysis, datasets were compared using the Kruskall-Wallace one-way analysis of PCR data was compared with unpaired two-sample t-tests. variance Error bars indicate standard deviation, and asterisks indicate statistical significance (^*^p<0.05, ^**^p<0.01, ^***^p<0.001, ^****^p<0.0001).

## Results

### Agarose-embedded macrophages form spheroids with long-term viability

To determine if macrophages alone can generate solid stress, we seeded single cells in an agarose hydrogel. As a plant-derived material, agarose is biologically inert to mammalian cells, and is also physically and chemically stable, with suitable biocompatibility [39, 40]. The mechanical properties of agarose are also easily tunable, as Young’s modulus increases exponentially with molar concentration [41]. Based on previous mechanical characterization by others and our own unconfined compression testing, we estimate that the Young’s moduli of 0.5%, 1%, and 2% agarose gels to be 2 kPa, 20 kPa, and 100 kPa, respectively [41]. These approximately correspond to the stiffness ranges measured in the brain, healthy heart, and fibrotic scar tissue, respectively [26, 42, 43]. Because the embedded macrophages cannot degrade the surrounding matrix, any cell growth must cause solid stress, as the cells must displace the elastic material in order to proliferate and form 3-D spheroids.

We observed that RAW264.7 murine macrophages readily form spheroids when embedded in agarose at concentrations between 0.5% and 2%. Gels with lower than 0.5% agarose content were excluded due to manipulation challenges. The majority of cells seeded in gels between 0.5% and 4% were still viable 24 hours after seeding, with the only significant difference being between 0.75% and 3% agarose, as quantified by calcein AM (i.e., live cells) and propidium iodide (i.e., dead cells) staining (**Fig. 1a**). However, gels with more than 2% agarose content failed to produce spheroids; thus 2% agarose was used as the maximum concentration condition. Agarose-embedded macrophage spheroids display excellent long-term viability, with live spheroids observed at least 16 days after seeding (**Fig. 1b**). The size and topography of these spheroids are sensitive to agarose gel concentration (**Fig. 1c**). This maximum time in culture was limited only by the over-proliferation of cells present on the gel surface or in the surrounding media, rather than those embedded within the gels, but it is likely these constructs could be maintained for much longer periods. Macrophages grown in 0.5% agarose grew notably faster than those in 1% agarose, which were correspondingly faster than those in 2% agarose. This effect was visible by eye as the aggregates became large enough to see over time. This was also apparent by the increased rate of media consumption in lower percentage gels.

**Fig. 1.**
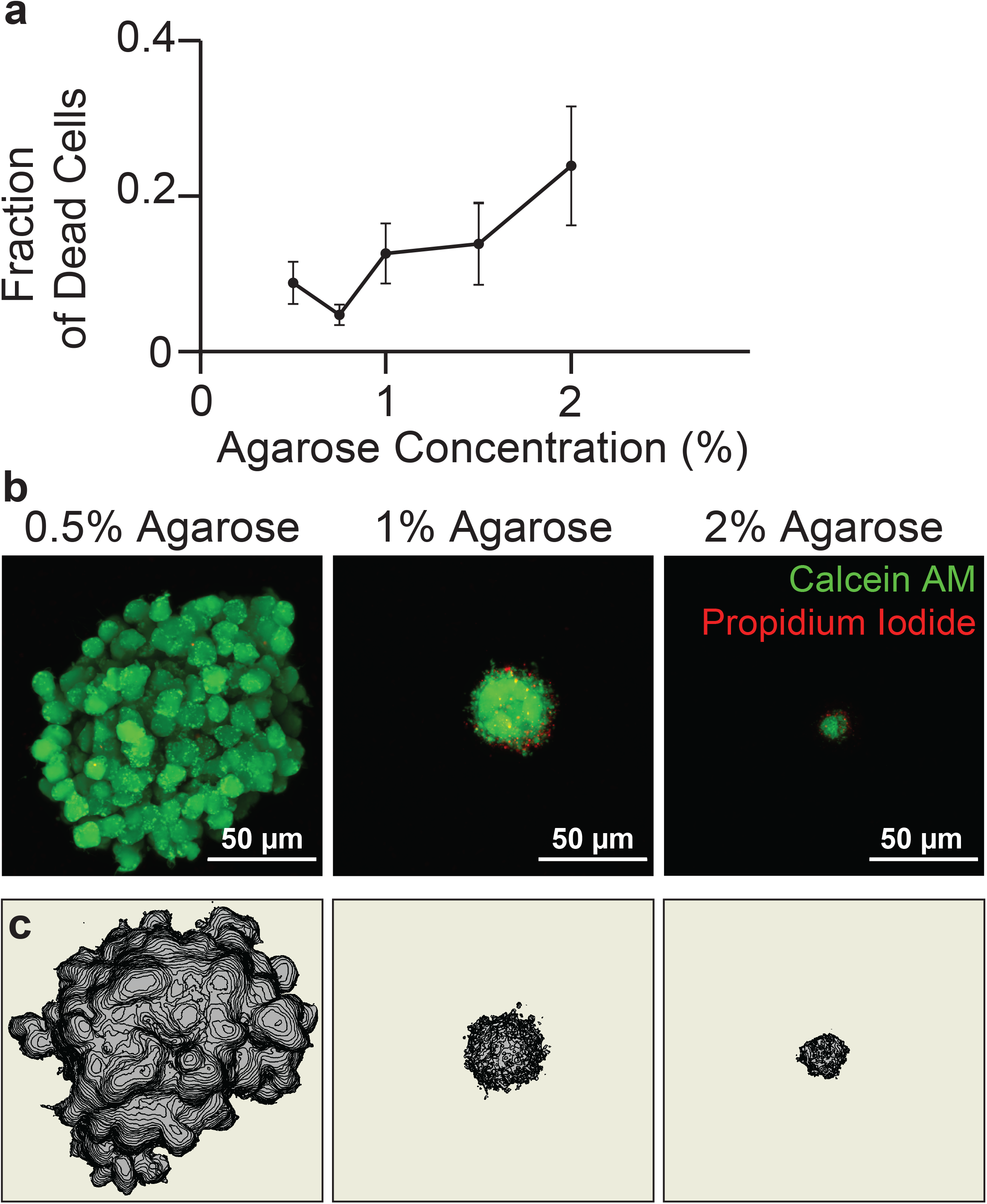
Macrophage spheroids embedded in agarose form aggregates with sustained viability. (**a**) Fraction dead cells 24h after seeding in agarose gels of varying concentration (**b**) Projections of macrophage spheroids 16 days after seeding show viable cells throughout the spheroid, in three different concentrations of agarose. Live cells are shown in green, dead cells shown in red, scale bar represents 50 µm. (**c**) Contour plots of the spheroid surface

### Macrophages generate solid stress in confining agarose gels

3-D imaging reveals that many spheroids adopt a flattened discoid shape, rather than a spherical shape (**Fig. 2**). Because macrophages cannot alter the plant-derived agarose matrix, any increase in spheroid size must cause displacement of the surrounding gel, thereby generating solid stress. Given an estimate of the spheroid geometry and the mechanical properties of the gel, we generated a simulation of the stress field in the gel surrounding the spheroids (**Fig. 2a**). Simulations of spheroid displacement during growth are shown based on fluorescent images. In **Fig. 2b** from left to right, the first three simulations for a spheroid embedded in 0.5%, 1% and 2% agarose gels assume no microcrack developments in the gels and an initial uniformly spherical geometry with a 5 μm radius, while the simulation for the spheroid in the 2% agarose gel with a discoid shape assumes formation and propagation of a microcrack in the gel during initial spheroid growth. The greatest solid stress-induced deformations are observed in 0.5% gels (with maximum gel displacements of ∼250 μm), compared to stiffer 2% gels with spherical (40 μm) and discoid (42 μm) shapes. However, solid stress propagates further into the surrounding matrix with increasing gel concentrations. These data demonstrate that increasing the rigidity of the surrounding agarose matrix by elevating gel concentration significantly restricts spheroid expansion and modulates growth morphology. The decreased gel displacement and altered shape in stiffer gels indicates that the increased mechanical resistance of the matrix impedes outward growth. In contrast, the more mechanically permissive and compliant 0.5% agarose gel allows for greater spheroid expansion which maintains a rounded shape.

**Fig. 2.**
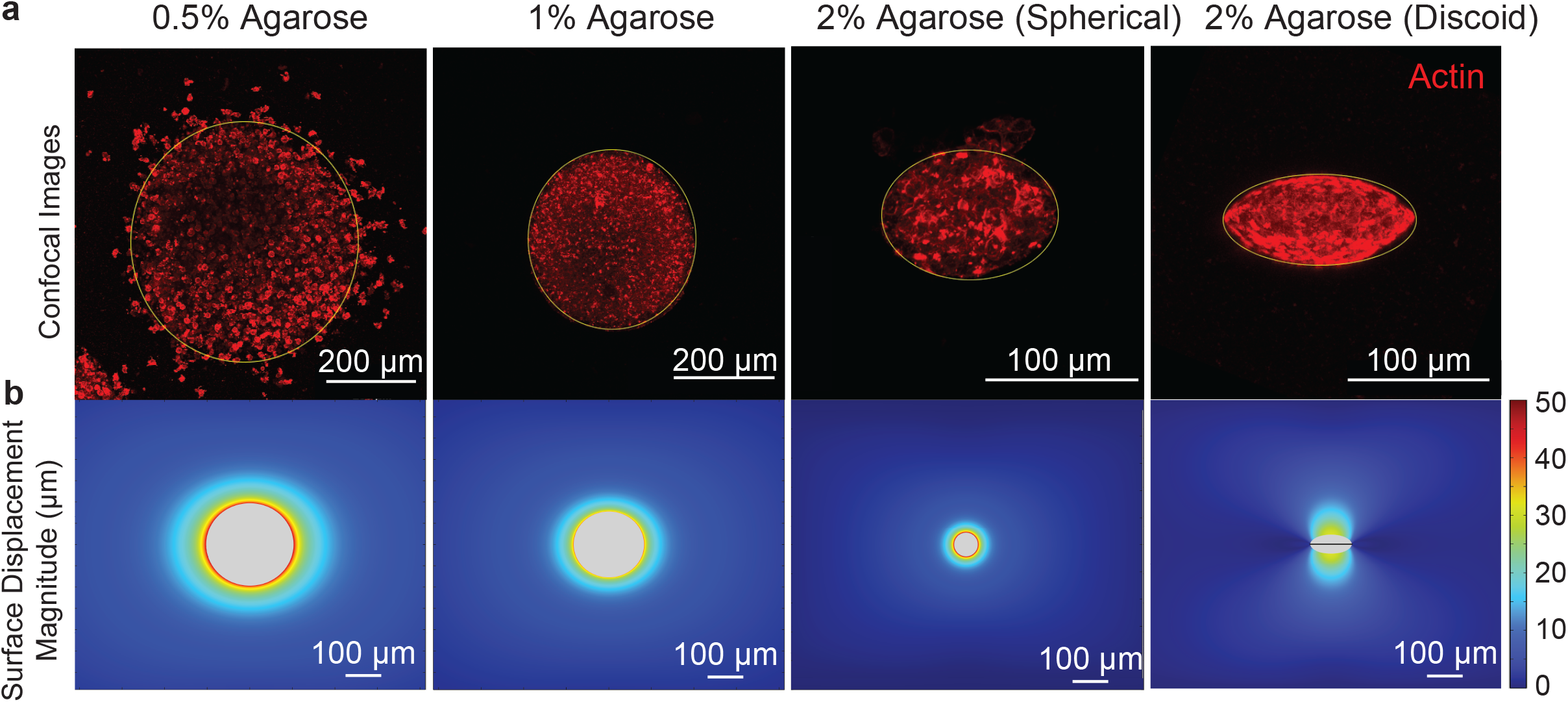
Agarose concentration modulates mechanical interactions and spheroid expansion. (**a**) Fluorescent imaging reveals that spheroids embedded in 0.5% agarose remain spherical, while those in 1% and 2% agarose become increasingly elongated and discoid. (**b**) Computational modeling visualizes the displacement field around expanding spheroids, with the greatest displacement seen in 0.5% agarose gels. Taken together, these results indicate that increasing agarose concentration restricts spheroid expansion and deformation, likely due to the increased mechanical rigidity restricting growth. The combined imaging and modeling approach provides visual evidence that agarose stiffness impacts spheroid morphology and expansion dynamics.

### Spheroid size decreases with increasing agarose concentration and increases with pro-inflammatory stimulation

Varying the concentration of agarose, as illustrated above, drastically impacts spheroid size without significantly altering viability within the range of 0.5% and 2% agarose, as determined by the ratio of propidium iodide-positive to calcein AM-positive cells. We also tested other biologically relevant stimuli, including addition of pro-inflammatory (M1-like) and anti-inflammatory (M2-like) polarizing cytokines, LPS stimulation, and hypoxic environments. For each of these conditions, we compared the spheroid area observed by brightfield microscopy. As agarose concentration decreases, the average spheroid size increases significantly (**Fig. 3a)**. In 0.5% agarose, spheroids are much less regular in shape than those in 1% and 2% as is apparent in images of fluorescently labeled actin at day 4 after seeding (**Fig. 3b**). Treating the agarose-embedded macrophages with 200 ng/ml LPS resulted in significantly increased spheroid size compared to untreated, for both 0.5% and 1% gels However, treatment with 20 ng/ml IFNγ in addition to 200 ng/ml LPS (i.e., M1-like polarization) decreased spheroid size. Treatment with 20 ng/ml of IL-4 (i.e., M2-like polarization) trended towards increased spheroid size. Interestingly, hypoxia did not significantly alter spheroid size.

**Fig. 3.**
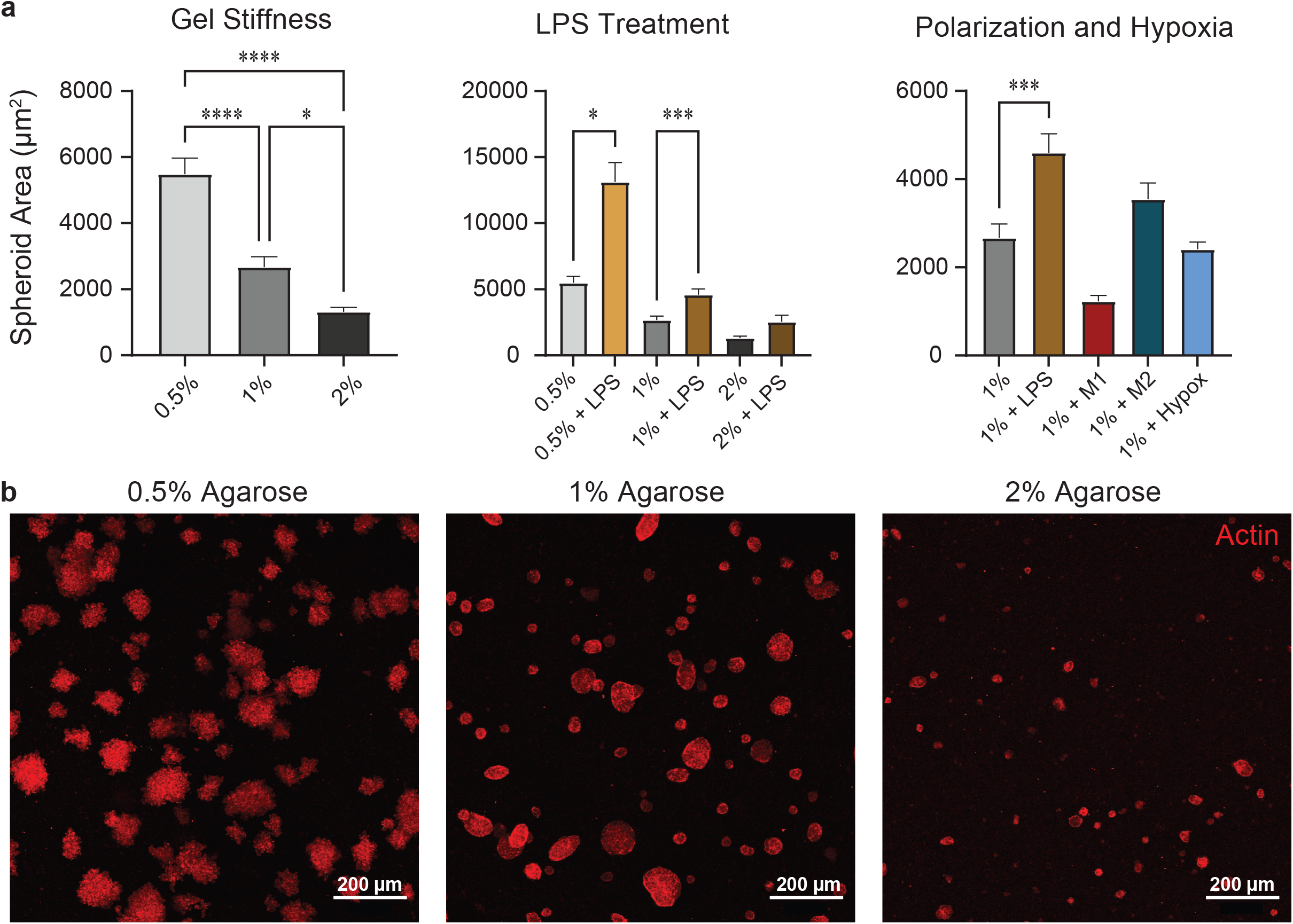
Spheroid size varies with agarose stiffness and macrophage stimulation/polarization. (**a**) Spheroid area, as quantified from brightfield images. (**b**) Representative images of maximum intensity projections of actin-stained spheroids (red)

### Stiffer gels reduce proliferation and mechanosensing protein expression in spheroids

As apparent in **Fig, 3b**, macrophage spheroids do not grow equally in all directions depending on their microenvironmental conditions, resulting in large-scale irregularity (e.g., ellipticity), and cell-scale differences in boundary uniformity (i.e., solidity). While we assumed in the simulation in **Fig. 2b** that discoid morphology is mechanically-driven (e.g., by micro-crack formation in the agarose gels), an alternative hypothesis could be tested that the irregular shape is biologically-driven (e.g., by anistropic proliferation). Therefore, we opted for a histological approach to capture any spatial differences in relevant biological markers that could be driving the asymmetry. Ki67 staining intensity decreased with increasing agarose concentration (**Fig. 4a**), corresponding to the decrease in spheroid size quantified above (**Fig. 3a**). The distribution of Ki67 cells throughout the spheroid did not clearly favor an edge versus central position (**Fig. 4b**). This indicates that differential rates of growth in different spatial directions may not be the result of variations in proliferation. This suggests that there may be a purely mechanical driving force behind the observed discoid shapes, such as the formation of cracks in the agarose gel and propagation of the spheroid into the crack space.

**Fig. 4.**
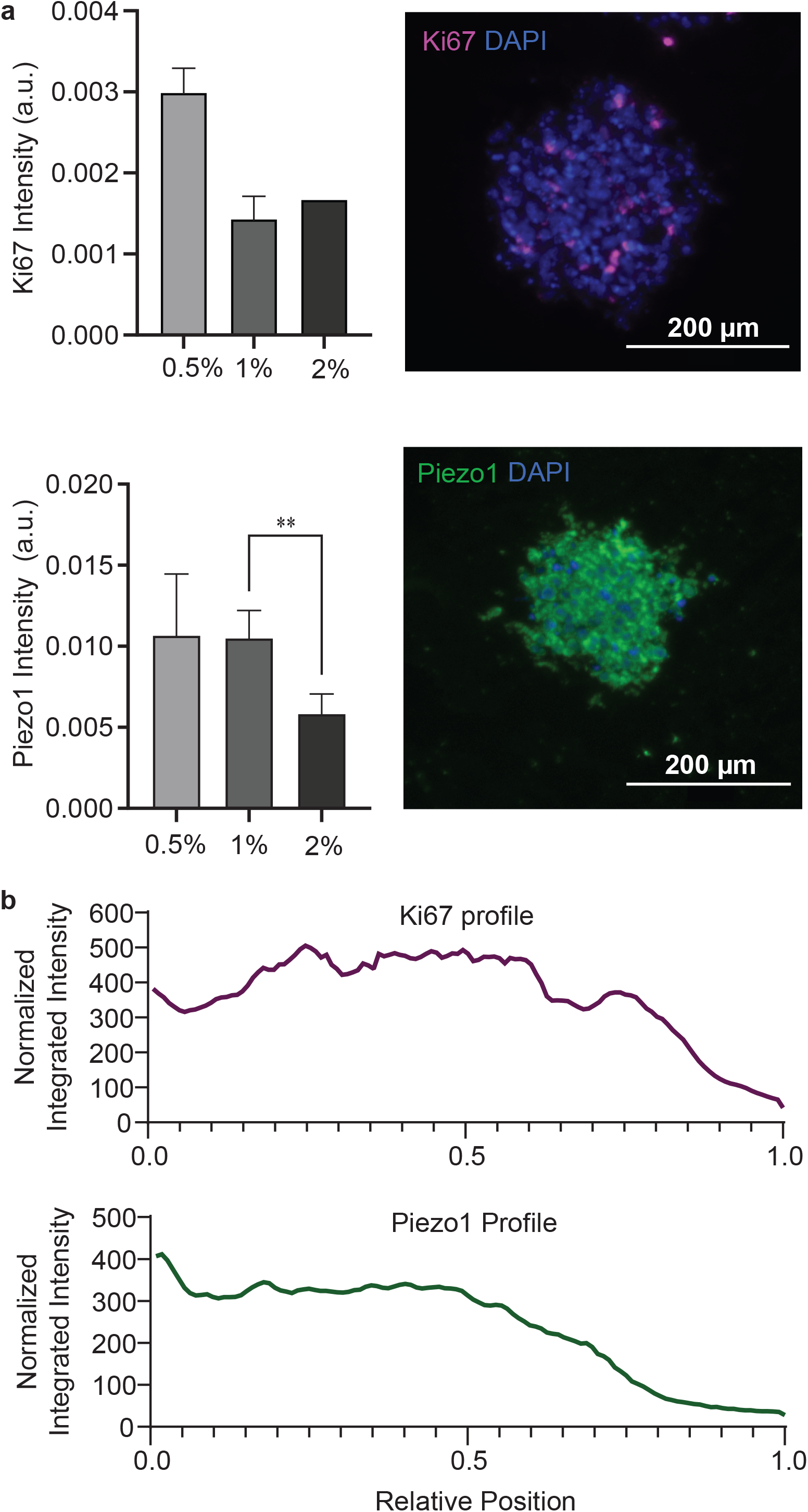
Immunofluorescent staining of spheroids sections for Ki67 and Piezo1. (**a**) Quantification and representative images of spheroid sections stained for Ki67(magenta) and Piezo1 (green). (**b**) Radial plots of the normalized integrated staining intensity of representative spheroids, with a relative position of 1 representing the spheroid edge

We also stained Piezo1, a mechanosensitive ion channel known to be involved in mechanotransduction in myeloid cells [44, 45]. Expression was distributed throughout the spheroid, with a slight peak at the center (**Fig. 4b**). In 2% agarose, Piezo1 expression was diminished, with a statistically significant decrease in average intensity between 1% and 2% agarose conditions (**Fig. 4a**). This result contrasts with prior studies showing that Piezo1 expression increases with stiffness in 2-D culture [46]. Piezo1 is a stretch-activated channel, but because mammalian cells cannot bind to the agarose, increased stiffness would not directly cause increased traction forces that are known to activate Piezo1 [47]. Piezo1 activation has been shown to promote myeloid cell expansion, so the concurrent increase in Piezo1 and Ki67 expression with reduced agarose concentration could be part of a causative relationship [44].

### Macrophages in 3D gels display altered mechanical and inflammatory transcriptional responses to varied mechano-chemical stimuli

We next compared the transcriptional activity of macrophages in various conditions. *Ki67* decreased significantly between 1% to 2% agarose (**Fig. 5a**). This aligns well with the spheroid size data (**Fig 3a**), implicating reduced proliferation behind the observed reduced spheroid size in stiffer gels. *Casp3*, an apoptosis marker, increased between 1% and 2%, but not between 0.5% and 1%. So, apoptosis is likely not responsible for the reduced size with increased stiffness, at least between 1% and 2% gels. Externally applied compression also decreased *Ki67* expression. M1-like polarization, M2-like polarization, and LPS stimulation all significantly decreased both *Ki67* and *Casp3* expression.

**Fig. 5.**
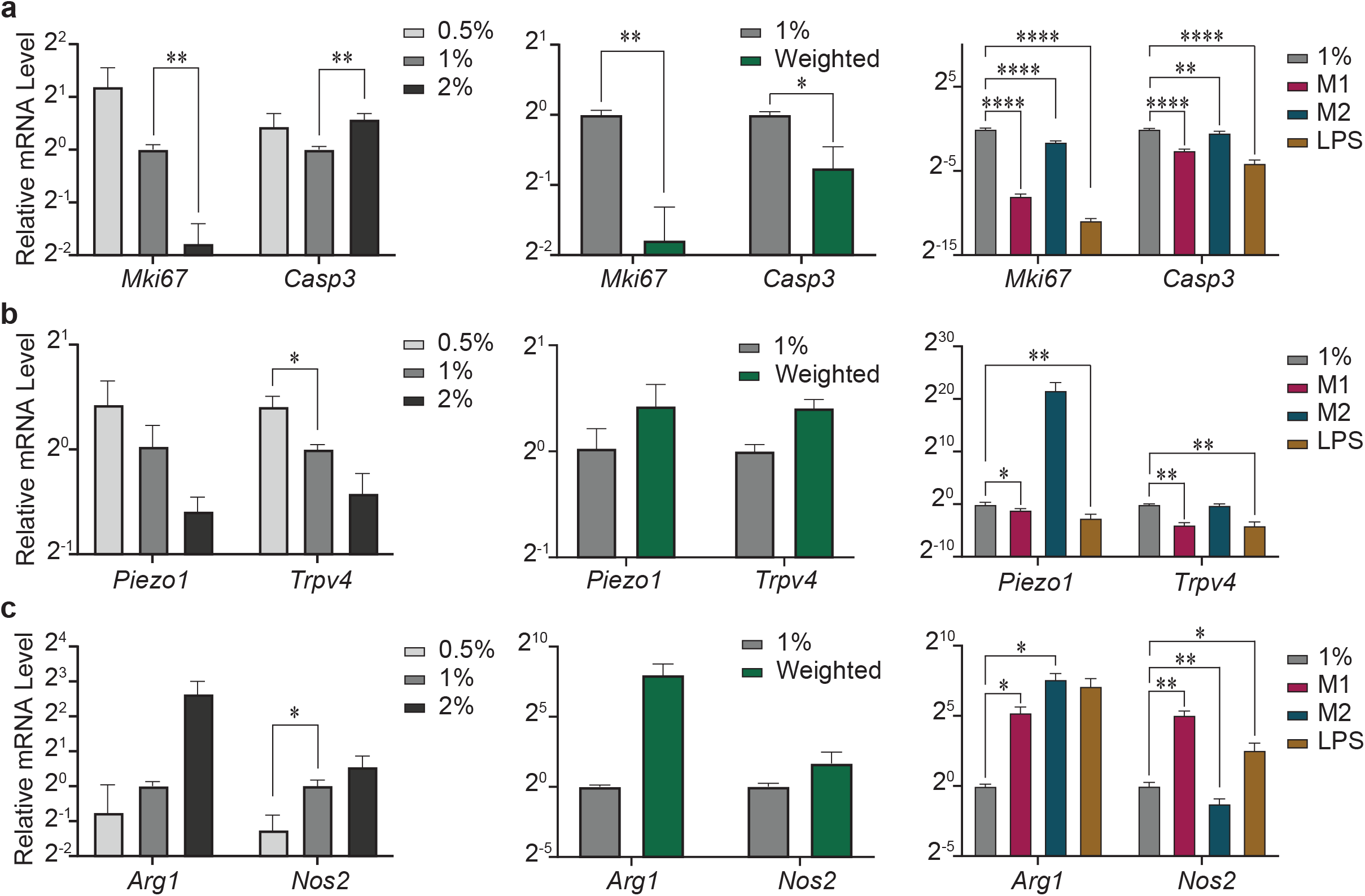
Macrophages show transcriptional response to both mechanical and chemical stimuli. qPCR analysis of genes related to proliferation and apoptosis (**a**), mechanosensitive ion channels (**b**), and macrophage polarization (**c**). Asterisks indicate comparisons which pass the false-discovery rate analysis

In addition to *Piezo1*, we also quantified *Trpv4*, another stretch-activated ion channel known to be involved in myeloid mechanotransduction [31, 48, 49]. *Trpv4* expression decreased between 0.5% and 1% agarose, and decreased with both M1 and LPS treatments (**Fig. 5b**). *Piezo1* expression also decreased with both M1 and LPS treatment. This potentially indicates that a decreased sensitivity to confining solid stress may play a role in the increase observed spheroid size between untreated and LPS-treated gels (**Fig. 3a**). Finally, we quantified canonical markers of pro-inflammatory macrophage activation (*Nos2*) and anti-inflammatory polarization (*Arg1*) (**Fig. 5c**). We observed a significant increase in *Nos2* with increasing gel stiffness as well as a trend towards increased *Arg1* with increasing gel stiffness. As expected, *Nos2* increased with M1 and LPS treatment, and decreased with M2 treatment. However, *Arg1* unexpectedly increased with all three stimulation/polarization treatments, potentially indicating altered phenotypic response to these stimuli in 3-D compared to previously observed 2-D culture results.

## Discussion

This 3-D model of macrophage spheroid formation and the accompanying computational modeling of stresses and strains and downstream cellular and molecular biology readouts on polarization combine to make a novel platform for investigating the immunomechanics and mechano-immunology of macrophages in varying biochemical and mechanical microenvironments. The agarose gel constructs are simple and inexpensive to create, making them attractive for high-throughput applications, such as drug screening. The model is also highly amenable to more complex co-culture or organoid experiments, as any number of cell types and treatments can be applied. It could therefore be used to model solid stress in other disease settings, such as within tumors or tuberculosis granulomas where macrophages dominate [50]. Additionally, the agarose gels can be processed identically to tissues for histological analysis. In this work, we cryopreserved the samples for immunofluorescent staining, but we have also successfully processed agarose-cell constructs for paraffin embedding and immunohistochemical staining. By increasing the density of cells initially seeded, we were also able to easily obtain sufficient RNA for multiple qPCR analyses.

To the best of our knowledge, this is the first demonstration of successful spheroid formation, long-term viability, and solid stress generation by macrophages alone. While many aspects of macrophage responses to various forms of mechanical stress have been studied, relatively little has been shown of the reciprocal regulation of macrophages and solid stress [7]. Our simulation results suggest that the mechanical microenvironment can override intrinsic growth programs to control spheroid expansion. Lower stiffness gels allow for a greater displacement of the gel around the spheroid compared to higher stiffness gels. However, in stiffer gels, the stress propagates further into the surrounding gel. This work also reveals an interesting 3-D growth pattern, as macrophages often adopt a flattened discoid shape, rather than a spheroidal shape. This could indicate either a physical process, such as the formation of a planar crack in the gel, or a biological process, such as differential proliferation or tip/leader cell migratory behavior in different regions of the spheroid [51]. Further work to characterize the agarose gel surrounding a spheroid and potential crack propagation is underway. A limitation of the computational model is that it provides an estimate of the displacement and solid stress of the agarose adjacent to the spheroid, but not within the spheroid itself. Future work to characterize the mechanical properties of macrophage spheroids would inform computational modeling of stress distribution within the spheroids.

The most potent biochemical cue that increases macrophage spheroid size and stress generation at all tested agarose concentrations is treatment with LPS, an immunostimulatory bacterial fragment used to mimic the effect of bacterial infection [52, 53]. Interestingly, the addition of IFNγ along with LPS, a standard M1-like polarizing regime, reversed the effect of LPS entirely. IFNγ and LPS are generally thought to synergize to induce an M1-like phenotype, but this 3-D stress-generation model seems to have identified a way in which the two oppose one another [54]. Further mechanistic studies utilizing this model will help to elucidate the nuances of LPS-stimulated versus canonically M1-like polarized macrophages. Prior studies have shown an inflammatory response when macrophages are cultured on stiffer 2-D substrates, but a more immunosuppressive phenotype when in a 3D matrix [27–34]. However, our results show an increase in an M1 marker with increased stiffness, indicating potential similarity with 2D observations.

In line with observations by others, TRPV4 expression decreased with stiffness as part of an M1-like response [31]. However, we also observed a large but not statistically significant increase in an M2 marker with stiffness. Piezo1 expression by macrophages has been shown to increase on stiffer surfaces [46, 55]. However, we observed a trend towards decreased Piezo expression with increased stiffness. Cells in a confining gel such as agarose that does not support cell adhesion may not experience the membrane tension that activates stretch-activated channels such as Piezo1 [56]. Further work will elucidate this 3D-specific mechanosensing response.

Macrophages have increasingly been a subject of interest as targets for disease treatments. A meta-analysis reports more than 25 clinical trials targeting tumor-associated macrophages in a range of different cancer types [57]. In atherosclerosis, reducing macrophage mass within plaques is a promising strategy, as is increasing the numbers of anti-inflammatory macrophages and decreasing pro-inflammatory macrophages in damaged heart tissue after myocardial infarction [18, 58]. Depleting macrophages in models of skin wounding reduces hypertrophic scar formation [19]. As previously shown, targeting either macrophage or solid stress in glioblastoma improves outcomes [59–61]. Thus, understanding factors that contribute to macrophage expansion or reduction via a novel 3-D system could inform future treatment strategies.

Overall, this work demonstrates a novel platform to study previously unexplored aspects of macrophage mechanics. Because many diseases involve both altered macrophage content and altered mechanics, this model may elucidate new and targetable pathological interactions between macrophages and solid stress. Understanding how macrophages generate stress, and how they respond to external cues under chronic solid stress, will inform strategies to target or reprogram macrophages in the plethora of diseases that involve macrophage accumulation. This platform also has promise for screening macrophage-targeted drugs and is highly tunable to apply to a range of diseases.

## Acknowledgments

Funding for this work was provided by the National Institutes of Health (NIH/NCI K22CA258410 and NIH/NIGMS R35GM151041), the Indiana Clinical Sciences and Translational Institute, and the Berthiaume Institute for Precision Health (all to M.D.). Other funding support included National Institutes of Health grants R01CA248033 (to X.L.) and R01CA280097 (to X.L., J.L.) and Department of Defense grants W81XWH2010312, W81XWH2010332, HT94252310010 and HT94252310613 (to X.L.). Histology was performed at the Notre Dame Histology Core, and multiphoton imaging was performed at the Notre Dame Integrated Imaging Facility, with the assistance of Dr. Sara Cole. The authors thank Ms. R’nld Rumbach for technical assistance, and Prof. Donny Hanjaya-Putra and his laboratory for imaging assistance.

## Author Contributions

M.D. and A.B. conceived of and designed this study, A.B. performed experiments and analysis of experimental data, S.S. performed computational analyses, data curation and image processing, A.B., S.S., J.L., X.L., M.D. contributed to data analysis and interpretation, A.B., S.S., and M.D. wrote the manuscript, and all authors reviewed and approved the final version.

